# Genomic and phylogenetic features of the *Picobirnaviridae* suggest microbial rather than animal hosts

**DOI:** 10.1101/2024.02.04.578841

**Authors:** Sabrina Sadiq, Edward C. Holmes, Jackie E. Mahar

## Abstract

The RNA virus family *Picobirnaviridae* has traditionally been associated with the gastrointestinal systems of terrestrial mammals and birds, with the majority of viruses detected in animal stool samples. Metatranscriptomic studies of vertebrates, invertebrates, microbial communities, and environmental samples have resulted in an enormous expansion of the genomic and phylogenetic diversity of this family. Yet picobirnaviruses remain poorly classified, with only one genus and three species formally ratified by the International Committee of Virus Taxonomy. Additionally, an inability to culture picobirnaviruses in a laboratory setting or isolate them in animal tissue samples, combined with the presence of bacterial genetic motifs in their genomes, suggests these viruses may represent RNA bacteriophage rather than being associated with animal infection. Utilising a data set of 2,286 picobirnaviruses sourced from mammals, birds, reptiles, fish, invertebrates, microbial communities, and environmental samples, we identified seven consistent phylogenetic clusters likely representing picobirnavirus genera that we tentatively name *Alpha-, Beta-, Gamma-, Delta-, Epsilon-, Zeta-,* and *Etapicobirnavirus*. A statistical analysis of topological congruence between virus-host phylogenies revealed more frequent cross-species transmission than any other RNA virus family. In addition, bacterial ribosomal binding site motifs were more enriched in picobirnavirus genomes than in the two groups of established RNA bacteriophage – the *Leviviricetes* and *Cystoviridae*. Overall, our findings support the hypothesis that the *Picobirnaviridae* have bacterial hosts and provide a lower-level taxonomic classification for this highly diverse and ubiquitous family of RNA viruses.

## INTRODUCTION

A combination of metatranscriptomic sequencing and data mining of the Sequence Read Archive (SRA) has greatly enhanced our knowledge of the scale, diversity, and composition of the RNA virosphere. As well as documenting huge numbers of novel viruses, this research program has provided new insights into previously known families of RNA viruses. For example, major changes to taxonomic organisations have been proposed based on updated phylogenetic trees of RNA viruses (Shi *et al*., 2016; Wolf *et al*., 2020; Sadiq *et al*., 2022), and host ranges have been extended when viruses are detected in organisms with which they are not typically associated (Urayama *et al*., 2016; Geoghegan *et al*., 2021; Bonny *et al*., 2021).

Members of the double-stranded RNA virus family *Picobirnaviridae* (order *Durnavirales*, class *Duplornaviricetes*, phylum *Pisuviricota*) have traditionally been considered opportunistic enteric pathogens of mammals and birds, typically detected in animal faecal samples (Malik *et al*., 2014), and were recently reported in invertebrate hosts (Delmas *et al*., 2019). In mammalian and avian species, picobirnavirus infection has been associated with disease presenting with diarrhoea or gastroenteritis (Malik *et al*., 2014; Ganesh *et al*., 2014). However, as picobirnaviruses have also been detected in the stool of healthy or asymptomatic animals (Ganesh *et al*., 2014), the presence of a picobirnavirus in the animal gastrointestinal system is not necessarily associated with overt disease. The genomes of these picobirnaviruses are typically bi-segmented, with segment 1 (2.4-2.7kb in length) comprising three open reading frames (ORF), although only the function of ORF3 is known, encoding a capsid protein precursor. Segment 2 (1.7-1.9kb) encodes the viral RNA-dependent RNA polymerase (RdRp) that is universal to RNA viruses and hence a powerful phylogenetic marker (Delmas *et al*., 2019). Picobirnaviruses with non-segmented genomes have also been reported, in which all ORFs are present in a single, monopartite genome (Giannitti *et al*., 2015; Shi *et al*., 2016).

The increasing accessibility of metagenomic sequencing has resulted in the identification of huge numbers of picobirnavirus sequences from both aquatic and terrestrial environmental samples such as wastewater, sewage, permafrost, farmland soils, and forest soils (Adriaenssens *et al*., 2018; Bell *et al*., 2020; Guajardo-Leiva *et al*., 2020; Chen *et al*., 2022; Neri *et al*., 2022). As metatranscriptomic sequencing necessarily produces a snapshot of all the RNA expressed in a sample, the host organisms for the majority of recently identified picobirnaviruses cannot be easily determined and characterisation of novel virus species largely relies on phylogenetic placement, often without the presence of complete virus genomes. While the number of picobirnavirus and picobirna-like sequences available on NCBI/GenBank is now in the thousands and comprises considerable genetic diversity, only a single genus and three species have been formally ratified in the most recent report of the International Committee on Taxonomy of Viruses (ICTV) (Delmas *et al*., 2019). The *Picobirnaviridae* are therefore in clear need of further taxonomic assessment and classification. An additional complicating factor is the large number of picobirnaviruses found within individual animal species has meant that multiple picobirnaviruses have been assigned the same name. For example, there are 12 viral RdRp sequences designated as ‘bovine picobirnavirus’, 13 as ‘dromedary picobirnavirus’, 16 as ‘human picobirnavirus’, and over 500 as ‘porcine picobirnavirus’ on NCBI/GenBank. Despite their identical names, these sequences are distinct and do not group together in phylogenetic trees (Delmas *et al*., 2019; Ghosh and Malik, 2021).

The remarkably wide range of both animal and non-animal sources in which picobirnaviruses have been detected, combined with an ongoing inability to propagate these viruses in eukaryotic cell lines or detect them in solid tissue samples, has led to the proposal that the *Picobirnaviridae* are not exclusively animal-associated, or that animals may not be the true hosts (Ghosh and Malik, 2021). The closest relative to the *Picobirnaviridae* is the plant-, fungi-, and protist-infecting family *Partitiviridae* (Vainio *et al*., 2018), members of which have also been detected in animal faeces (Chen *et al*., 2021) and invertebrate samples (Shi *et al*., 2016; Le Lay *et al*., 2020), suggesting they infect components of animal diet rather than the animal themselves. Hence, it is reasonable to propose that the *Picobirnaviridae* are similarly associated with animal diet and/or commensal gut microflora. Aspects of picobirnavirus genomes also support the hypothesis that these viruses do not infect animals (at least not exclusively) and may instead be associated with microbial organisms. In particular, the prokaryotic ribosomal binding site (RBS) motif, necessary for the initiation of translation in prokaryotes, is present in the genomes of other RNA bacteriophage such as in the *Cystoviridae* (Ghosh and Malik, 2021). A high prevalence of RBS motif sequences has been identified upstream of ORFs in both picobirnavirus genome segments (Krishnamurthy and Wang, 2018), suggesting that the picobirnaviruses in fact predominantly represent a family of bacteriophage. In addition, several picobirnavirus species identified from mammals and invertebrates (Yinda *et al*., 2019; Kleymann *et al*., 2020) utilise the mitochondrial genetic code, a characteristic shared by members of the RNA virus family *Mitoviridae* (phylum *Lenarviricota*) that replicate in the fungal mitochondria (Cole *et al*., 2000; Hillman and Cai, 2013).

Using a set of 2,286 publicly available *Picobirnaviridae* RdRp sequences, the majority of which were generated through metatranscriptomic sequencing, we aimed to provide a realistic classification of this important group of viruses, identifying distinct genera, and to reassess their host range. We based our analysis on the extent of congruence between virus and host phylogenetic trees, as well as on features of *Picobirnaviridae* genomes that may be indicative of bacterial hosts.

## METHODS

### Data set collection and processing for phylogenetic analysis

For all phylogenies generated in this work, the number of sequences included in the data set, the outgroups used, and the lengths (trimmed and untrimmed) of each alignment generated are detailed in Table 1. The labels T1 through T21, where ‘T’ simply stands for ‘tree’, refer to an alignment or phylogeny generated on a particular combination of *Picobirnaviridae* and outgroup family sequences. The exception is T21, which is a host animal cladogram artificially generated in Newick format and used to assess host associations. The data sets assembled for each tree are detailed below.

**Table 1.** Details of sequence data sets, alignments and rooting schemes used to generate phylogenetic trees.

| Tree | Sequences analysed (total number) | Alignment length (amino acid residues) |  | Rooting scheme | Figure(s) |
| --- | --- | --- | --- | --- | --- |
|  |  | Full length | Trimmed |  |  |
| **T1** | *Picobirnaviridae* (2,286) | 13,540 | 704 | Midpoint | Fig. 1A |
| **T2** | *Picobirnaviridae* (2,286),<br>*Amalgaviridae* (14) | 13,430 | 712 | *Amalgaviridae* | Fig. 1B |
| **T3** | *Picobirnaviridae* (2,286),<br>*Curvulaviridae* (8) | 13,744 | 701 | *Curvulaviridae* | Fig. 1C |
| **T4** | *Picobirnaviridae* (2,286),<br>*Fusariviridae* (32) | 13,738 | 701 | *Fusariviridae* | Fig. 1D |
| **T5** | *Picobirnaviridae* (2,286),<br>*Partitiviridae* (2,536) | 18,017 | 757 | *Partitiviridae* | Fig. 1E |
| **T6** | *Picobirnaviridae* (2,286),<br>*Partitiviridae* (2,536),<br>*Amalgaviridae* (14) | 17,740 | 710 | *Amalgaviridae* | Fig. 1F |
| **T7** | *Picobirnaviridae* (2,286),<br>*Partitiviridae* (2,536),<br>*Curvulaviridae* (8) | 17,742 | 710 | *Curvulaviridae* | Fig. 1G |
| **T8** | *Picobirnaviridae* (2,286),<br>*Partitiviridae* (2,536),<br>*Fusariviridae* (32) | 17,231 | 706 | *Fusariviridae* | Fig. 1H |
| **T9** | *Picobirnaviridae* (2,286),<br>*Partitiviridae* (40) | 13,906 | 695 | *Partitiviridae* | Fig. 1I, Fig.<br>4B, D, Supp.<br>Fig. 9, 11 |
| **T10** | *Picobirnaviridae* (2,286),<br>*Partitiviridae* (394) | 14,063 | 703 | *Partitiviridae* | Fig. 1J |
| **T11** | *Durnavirales* excluding<br>family *Hypoviridae*<br>(4,717) | 14,971 | 808 | Unrooted | Fig. 2 |
| **T12** | *Alphapicobirnavirus*<br>(421) | 3,639 | 728 | Midpoint | Fig. 3A,<br>Supp. Fig. 1 |
| **T13** | *Betapicobirnavirus* (802) | 2,844 | 711 | Midpoint | Fig. 3B,<br>Supp. Fig. 2 |
| **T14** | *Gammapicobirnavirus*<br>(418) | 3,856 | 733 | Midpoint | Fig. 3E,<br>Supp. Fig. 5 |
| **T15** | *Deltapicobirnavirus*<br>(311) | 5,759 | 749 | Midpoint | Fig. 3F,<br>Supp. Fig. 6 |
| **T16** | *Epsilonpicobirnavirus*<br>(36) | 853 | 691 | Midpoint | Fig. 3C,<br>Supp. Fig. 3 |
| **T17** | *Zetapicobirnavirus* (67) | 824 | 700 | Midpoint | Fig. 3D,<br>Supp. Fig. 4 |
| **T18** | *Etapicobirnavirus* (70) | 881 | 705 | Midpoint | Fig. 3G,<br>Supp. Fig. 7 |
| **T19** | Animal-associated<br>*Picobirnaviridae* (1,269),<br>*Partitiviridae* (40) | 5,734 | 745 | *Partitiviridae* | Fig. 4A, C,<br>Supp. Figs.<br>8, 10 |
| **T20** | Animal-associated<br>*Picobirnaviridae* (1,269) | 5,591 | 727 | Midpoint | Input for<br>Fig. 5 (not<br>shown) |
| **T21** | NA – artificially<br>generated host cladogram | NA | NA | NA | Input for<br>Fig. 5,<br>Supp. Fig. 12 |

As a base *Picornaviridae* data set, curated in June 2022, we collated all non-redundant picobirnavirus RdRp sequences available on NCBI/GenBank by searching the protein database for “picobirna”. Only sequences with associated hosts or sampling environment metadata were included. Picobirnavirus and picobirna-like sequences described by Neri *et al*. (2022) were also added to this data set. The majority (89%) of sequences collected were faecal or environmentally sourced picobirnaviruses described in Chen *et al*. (2022) and Neri *et al*. (2022). A total of 3,644 sequences clustered at 90-99% amino acid identity using CD-HIT (Fu *et al*., 2012). One representative sequence from each cluster was retained. The total number of picobirnavirus sequences collated in this manner for further analysis was 2,286.

RdRp sequences from families related to the *Picobirnaviridae* within the order *Durnavirales* were included in additional alignments as outgroups to provide directionality to the resulting phylogenies: the *Amalgaviridae* (14 sequences; T2, T6 in Table 1), *Curvulaviridae* (8 sequences; T3, T7), *Fusariviridae* (32 sequences; T4, T8), and the *Partitiviridae* (40, 394, or 2,536 sequences; T9, T10, and T5-8, respectively). A phylogeny of the order *Durnavirales* was estimated on 4,717 sequences from the families *Amalgaviridae, Curvulaviridae, Fusariviridae, Picobirnaviridae*, and *Partitiviridae* (T11). Phylogenetic trees were also estimated based on sequence alignments of each of the proposed *Picobirnaviridae* genera (T12-18). Finally, sequence alignments were constructed on a subset of 1,269 picobirnavirus sequences detected in animal (typically faecal) samples, with (T19) and without (T20) an outgroup of 40 partitivirus sequences.

### Sequence alignment and phylogenetic analysis

All sequences were aligned using MAFFT (v7.487) (Katoh and Standley, 2013) and trimmed using trimAL (v1.4.1) (Capella-Gutierrez *et al*., 2009) to retain the most conserved amino acid positions. Trimmed alignments were manually inspected in Geneious Prime (v.11.0.14.1) to identify and remove any ambiguously aligned regions, resulting in trimmed alignments ranging between 691-757 positions in length used to generate phylogenetic trees.

The best-fit amino acid substitution model for all data sets was found to be the Le-Gascuel (LG) model, established using ModelFinder within IQ-TREE (v1.6.2). Due to the consistently very high number of amino acid changes per position, a gamma distribution of among-site rate variation was not used to estimate the full *Durnavirales* phylogeny (T11 in Table 1). Maximum likelihood phylogenetic trees were then estimated on each data set using IQ-TREE (v1.6.2) (Nguyen *et al*., 2015), with node support evaluated using the SH-like approximate likelihood ratio test (SH-aLRT), with 1,000 replicates. Trees were visualised in FigTree (v1.4.4) (Rambaut, 2018), as well as in R (v4.1.0) using the packages ‘ape’ (v5.5) (Paradis and Schliep, 2019) and ggtree (v3.0.2) (Yu *et al*., 2017).

### Classification of the *Picobirnaviridae*

Ten phylogenies (T1-T10; see Table 1) were used for determining potential genera within the *Picobirnaviridae*. Four large and three small defined clusters were identified in the midpoint-rooted tree (T1 in Table 1), defined by relatively long branch lengths to their respective common ancestor node and ≥80% SH-aLRT support. Sequences within each cluster were annotated as ‘clade 1’ through ‘clade 7’. These clusters, or “draft genera’’ were then annotated in all other trees. Any sequences not clustering within their assigned “draft genus” with ≥80% SH-aLRT node support in any of the ten trees were excluded. The remaining sequences that consistently grouped within the same clade in every tree were considered the “core” sequences of each proposed genus. Sequence alignments were constructed on each set of core sequences and subsequent genus-level phylogenetic trees were estimated as described above.

### Analysis of phylogenetic incongruence

Prior to analysis, *Picobirnaviridae* sequences were grouped by assigned host (animals) or sampling source (environments and microbial communities). Hosts and sampling sources were categorised in two ways (Table 2). The first focused on specific animal hosts, with non-animal sources simply reflected in a joint ‘microbial/environmental’ category, while mammals were further categorised into lower taxonomic levels. The second grouped animal hosts more broadly as ‘mammalian’, ‘avian’, ‘fish’, ‘reptile’, and ‘invertebrate’, with non-animal sources similarly expanded into ‘aquatic’, ‘terrestrial’, and ‘engineered’ environments and microbial communities. Aquatic sources included bodies of water and sediments, while terrestrial environments included various soils and plant matter (Chen *et al*., 2022; Neri *et al*., 2022). Engineered sources included wastewater, food fermentation processes, and laboratory cultures (Neri *et al*., 2022).

**Table 2.** Categories of assumed host animal or environmental/microbial source of the *Picobirnaviridae* sequences analysed in this study.

| Animal-host focused list | Environmental/microbial source-focused list |
| --- | --- |
| Primate | Mammalian |
| Bovidae |  |
| Cervidae |  |
| Equidae |  |
| Suidae (porcine) |  |
| Dromedary |  |
| Feline |  |
| Canine |  |
| Tasmanian devil |  |
| Rodent |  |
| Lagomorph |  |
| Bat |  |
| Avian | Avian |
| Reptile | Reptile |
| Fish | Fish |
| Arthropod | Invertebrate |
| Other invertebrate |  |
| Microbial/environmental | Microbial - aquatic |
|  | Microbial - engineered |
|  | Microbial - terrestrial |
|  | Environmental - aquatic |
|  | Environmental - engineered |
|  | Environmental - terrestrial |

A cladogram of relevant ‘host’ animals (as listed in Table 2) was constructed based on the evolutionary relationships demonstrated in the current literature on vertebrate and invertebrate evolution (Lindblad-Toh *et al*., 2005; Song *et al*., 2012; Fong *et al*., 2012; Shen *et al*., 2014; Prum *et al*., 2015; Zhou *et al*., 2017; Zurano *et al*., 2019). The eMPRess program (v1.2.1) (Santichaivekin *et al*., 2021) was used to reconcile topologies of the animal-associated picobirnavirus (T20 in Table 1) and host animal (T21 in Table 1) phylogenies to determine the event likelihoods of co-divergence, cross-species transmission, duplication, and extinction. Event costs were defined as 0 for co-divergence and 1 for cross-species transmission, duplication, and extinction. The R package ‘NELSI’ (v0.21) (Ho *et al*., 2015) was used to calculate the normalised PH85 (nPH85) distance for the *Picobirnaviridae*. The nPH85 distance is based on Penny and Hendy distance metric and describes the topological distance between the virus and host phylogenies (Geoghegan *et al*., 2017). An nPH85 distance of 0 indicates complete co-divergence between virus and host, whereas an nPH85 of 1 indicates complete cross-species transmission.

### Presence of bacterial ribosomal binding sites

A sequence alignment was generated using MAFFT (v7.487) comprising all available non-redundant and annotated nucleotide sequences of segment 1 of the picobirnavirus genome that contained at least 30 nucleotides of a 5’ untranslated region (UTR), totalling 360 sequences. Sequences were aligned to assist in locating the start codons of ORFs, upstream of which the ribosomal binding site motifs are potentially located. Similarly, an alignment was constructed comprising all available segment 2 *Picobirnaviridae* sequences using the same criteria, for a total of 984 sequences. Occurrences of the ‘AGGAGG’ hexamer within 14-24 nucleotides upstream of each ORF’s start codon were identified and annotated using Geneious Prime (v11.0.14.1), as these likely represented a bacterial RBS motif and hence are evidence of a bacterial host. The frequency of RBS occurrence for each segment was calculated as a proportion of the total number of annotated ORFs in the set of sequences for that segment. To compare the frequencies of RBS presence between the *Picobirnaviridae* and established RNA bacteriophage families this process was repeated on 1,040 *Leviviricetes* genomes and 17, 11, and 38 *Cystoviridae* S, M, and L segments, respectively.

## RESULTS

### Classification of the *Picobirnaviridae*

We observed seven major well-supported clusters of RdRp sequences within the *Picobirnaviridae* that were consistently present across ten phylogenies inferred using different outgroups and rooting methods (Figure 1, Table 1). Each of the seven clades were defined by relatively long branches to their respective common ancestral node with >80% SH-aLRT node support in each of the ten phylogenies (Figure 1). We propose that these groups are candidates for novel genera with the *Picobirnaviridae*, tentatively named: *Alphapicobirnavirus* (421 sequences, containing *Orthopicobirnavirus equi*)*, Betapicobirnavirus* (802 sequences, containing *Orthopicobirnavirus hominis*)*, Gammapicobirnavirus* (418 sequences, containing *Orthopicobirnavirus beihaiense*)*, Deltapicobirnavirus* (311 sequences), *Epsilonpicobirnavirus* (36 sequences), *Zetapicobirnavirus* (67 sequences), and *Etapicobirnavirus* (70 sequences), in line with the nomenclature for other multi-genus families within the *Durnavirales*. Sequences consistently clustering within one of these clades in each of the ten trees were considered the “core” members comprising each genus. Those sequences whose phylogenetic position was inconsistent remained as “unclassified” *Picobirnaviridae*. Using this approach, we were able to assign 2,125 non-redundant picobirnaviruses to a proposed genus (Supplementary Table 1). Genus-level phylogenies of each proposed genus are shown in Supplementary Figures 1-7.

**Figure 1.**
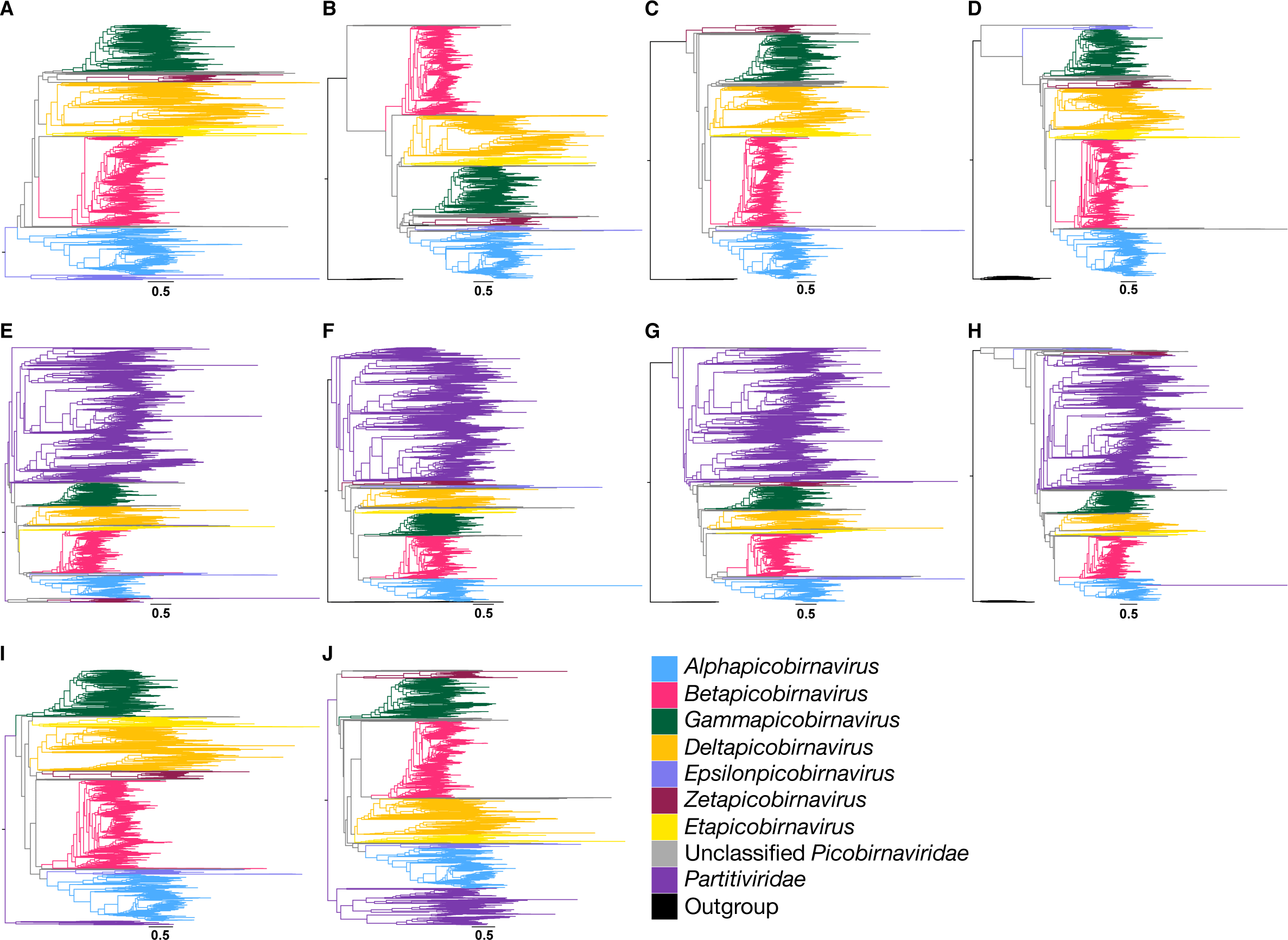
Maximum likelihood phylogenies of the family *Picobirnaviridae* estimated on the RdRp and using different families of the order *Durnavirales* as outgroups. Row one contains the *Picobirnaviridae*: (A) midpoint-rooted or utilising the (B) *Amalgaviridae*, (C) *Curvulaviridae*, or (D) *Fusariviridae* as outgroups. Row two contains the multi-family *Partitivirdae-Picobirnaviridae* clade: (E) midpoint-rooted or utilising the (F) *Amalgaviridae*, (G) *Curvulaviridae*, or (H) *Fusariviridae* as outgroups. Row three contains the *Picobirnaviridae* with a subset of (I) 40 or (J) 394 *Partitiviridae* sequences as outgroups. In all trees, the *Partitiviridae* are marked by purple branches and the outgroup is in black. The proposed *Picobirnaviridae* genera – *Alpha*-, *Beta*-, *Gamma*-, *Delta*-, *Epsilon*-, *Zeta*-, and *Etapicobirnavirus* – are coloured by blue, pink, green, orange, light purple, maroon, and yellow branches, respectively, while sequences unable to be assigned to a genus are represented by grey branches. Horizontal branches are drawn to scale, and the scale bars below each tree represent 0.5 amino acid substitutions per site.

The only currently accepted genus in the *Picobirnaviridae* – *Orthopicobirnavirus* – contains three species: *Orthopicobirnavirus hominis, Orthopicobirnavirus equi,* and *Orthopicobirnavirus beihaiense*, isolated from human (Wakuda *et al*., 2005), horse (Giannitti *et al*., 2015), and peanut worms (Shi *et al*., 2016), respectively. Each of these species appear in a different proposed genus described here (Supplementary Figures 1-3), limiting our ability to clearly determine which most closely resembles the genus *Orthopicobirnavirus*.

There were no common topologies for the proposed *Picobirnaviridae* genera across all ten phylogenetic trees. As a consequence, exact evolutionary relationships among genera cannot be safely determined. However, the genera *Deltapicobirnavirus* and *Etapicobirnavirus* formed sister clades in seven of the ten phylogenies (Figure 1A-D, F, H, J). In the remaining three trees (Figure 1E, G, I), one genus fell directly basal to the other, but which genus was basal to the other was inconsistent among trees. In six phylogenies, *Alphapicobirnavirus* and *Epsilonpicobirnavirus* formed sister clades (Figure 1B, C, E, G, I, J), suggesting that these genera are more closely related to each other. In eight trees, the two largest genera, *Alphapicobirnavirus* and *Betapicobirnavirus,* either grouped together (Figure 1D-I) or one fell directly basal to the other (Figure 1A, C). Furthermore, in four of the trees where *Alphapicobirnavirus* and *Betapicobirnavirus* genera clustered together, *Epsilonpicobirnavirus* was also part of the group as a sister clade to *Alphapicobirnavirus* (Figure 1C, E, G, I).

We next sought to give these proposed genera evolutionary context by placing them in a tree with multiple related families of RNA viruses. An unrooted phylogeny was estimated on 4,717 RdRp sequences from the order *Durnavirales*, including the *Picobirnaviridae*, *Partitiviridae*, *Amalgaviridae*, *Curvulaviridae*, and *Fusariviridae* (Figure 2). Sequences from the family *Hypoviridae* were too divergent to reliably align and were therefore excluded. Importantly, the *Picobirnaviridae* formed a monophyletic group that was phylogenetically distinct from the other families in the order. Certain trends in tree topology frequently observed in Figure 1 were repeated here, such as *Deltapicobirnavirus* and *Etapicobirnavirus* forming sister clades, as well as *Alphapicobirnavirus* and *Epsilonpicobirnavirus* forming sister clades that clustered with *Betapicobirnavirus* (Figure 2). The proposed genera also remained mostly intact despite the high level of diversity in the alignment. The only exception was a group of seven divergent deltapicobirnaviruses that fell as a small sister clade to the zetapicobirnaviruses (Figure 2, denoted by a star).

**Figure 2.**
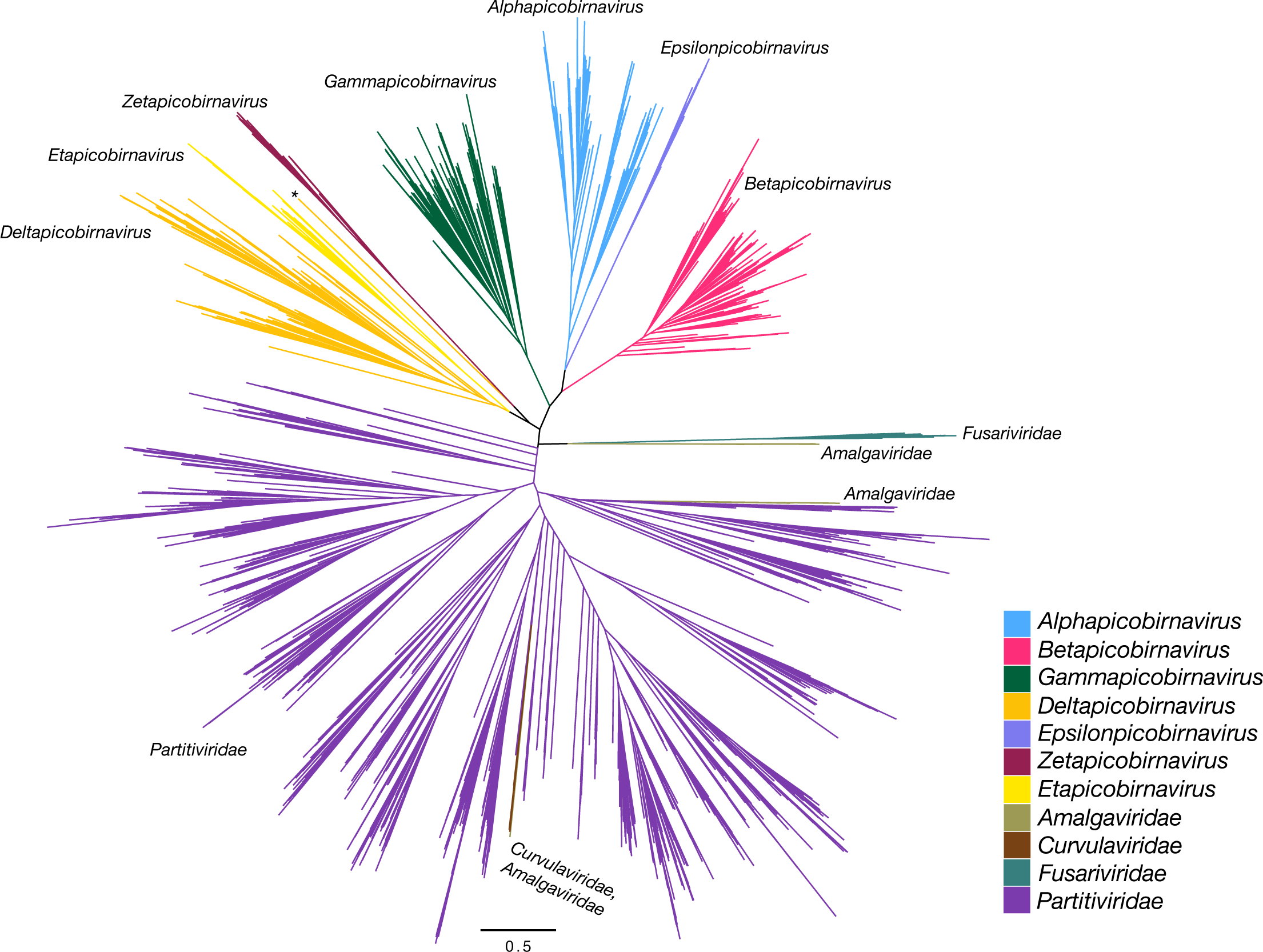
Unrooted maximum likelihood tree of 4,717 *Durnavirales* RdRp sequences, with branches coloured by family (*Amalgaviridae*, *Curvulaviridae*, *Fusariviridae*, *Partitiviridae*) or proposed genus (*Picobirnaviridae*). Families *Amalgaviridae*, *Curvulaviridae*, *Fusariviridae*, and *Partitiviridae* are marked by olive-green, brown, teal, and purple branches, respectively. The proposed *Picobirnaviridae* genera – *Alpha*-, *Beta*-, *Gamma*-, *Delta*-, *Epsilon*-, *Zeta*-, and *Etapicobirnavirus* – are coloured by blue, pink, green, orange, light purple, maroon, and yellow branches, respectively, while sequences unable to be assigned to a *Picobirnaviridae* genus are represented by grey branches. Branch lengths indicate the number of amino acid substitutions per site, as represented by the scale bar.

### Distribution of apparent hosts or environmental source of picobirnaviruses in each proposed genus

The genus *Alphapicobirnavirus* (Figure 3A, Supplementary Figure 1) predominantly comprised mammalian and avian picobirnaviruses, as well as those sampled from some invertebrate species. In the most basal clade of this genus the majority of sequences were identified in microbial communities, although a small number of mammalian, avian, invertebrate, and environmentally-sourced picobirnaviruses also fell in this group. The genus *Betapicobirnavirus* (Figure 3B, Supplementary Figure 2) was similarly dominated by picobirnaviruses from mammalian and avian sources. This genus also contained the divergent species identified in fish (AVM87403 Beihai goldsaddle goldfish picobirnavirus) and lizards (AVM87436 Guangdong Chinese water skink picobirnavirus and UCS96434 Picobirnaviridae sp.). There were fewer microbial/environmental-sourced viruses in the betapicobirnaviruses and none formed large, monophyletic clusters as seen in the basal *Alphapicobirnavirus* group.

**Figure 3.**
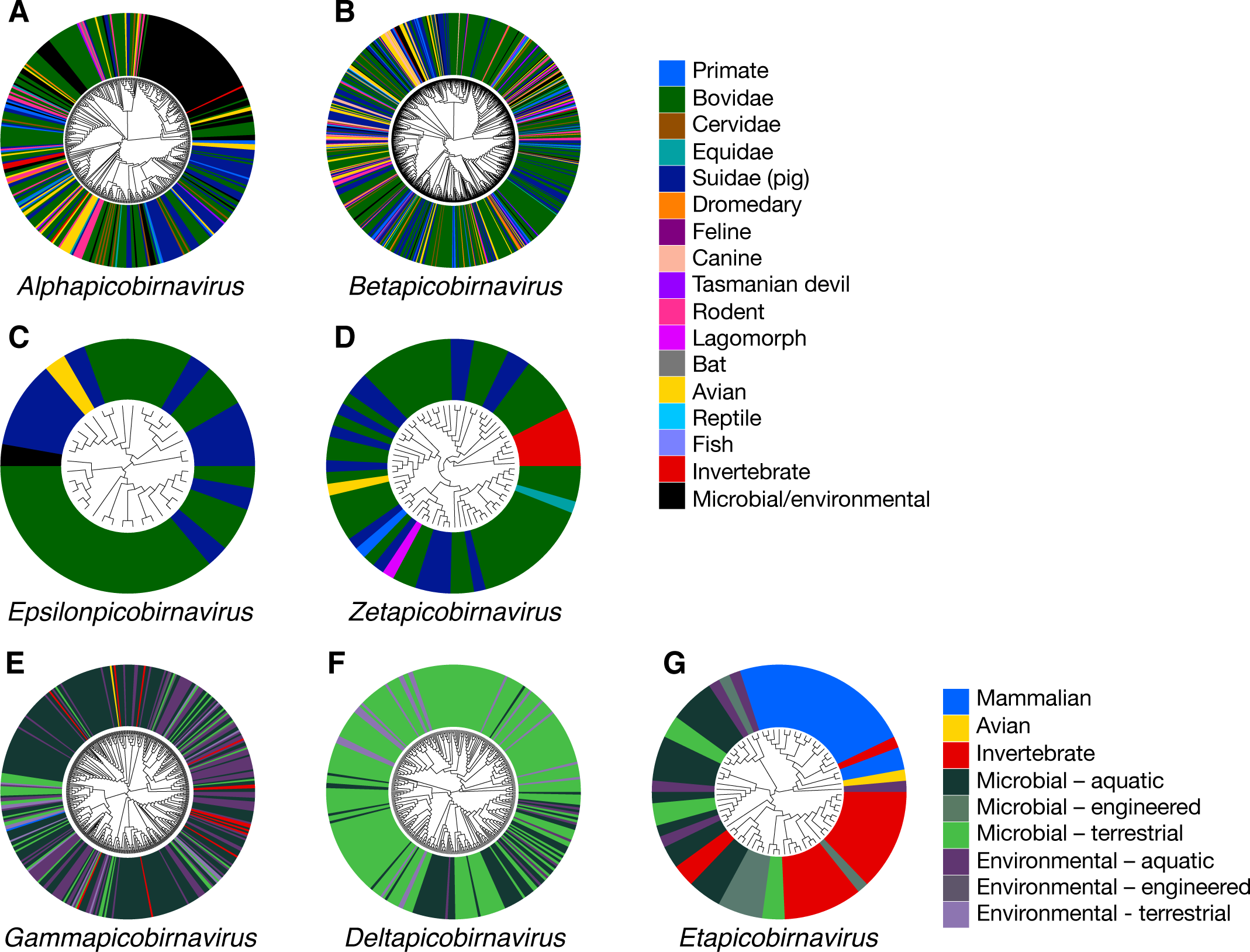
Maximum likelihood phylogenetic trees of the RdRp of proposed *Picorbirnaviridae* genera coloured by apparent host or sampling environment. All phylogenies are midpoint-rooted. Tip labels are coloured to represent assigned host/sampling environment. *Alphapicobirnavirus* (A), *Betapicobirnavirus* (B), *Epsilonpicobirnavirus* (C) and *Zetapicobirnavirus* (D) are coloured by specific assigned animal host. *Gammapicobirnavirus* (E), *Deltapicobirnavirus* (F), and *Etapicobirnavirus* (G) categorise animal hosts more broadly as ‘mammalian’, ‘avian’, and ‘invertebrate’, and non-animal associated sequences are grouped into ‘environmental’ or ‘microbial’ samples, both further classed into ‘terrestrial’, ‘aquatic’, and ‘engineered’ sampling sources. Branch lengths indicate the number of amino acid substitutions per site. Scale bars for branch lengths are shown in rectangular versions of trees (Supplementary Figures 1-7).

In contrast to the predominantly animal-associated genera described above, the *Gammapicobirnavirus* (Figure 3E, Supplementary Figure 3) and *Deltapicobirnavirus* (Figure 3F, Supplementary Figure 4) genera were almost entirely composed of sequences detected in microbial communities and environmental samples. Some invertebrate-associated picobirnaviruses (namely, from Cnidara, Porifera, crustaceans, and molluscs) clustered within the *Gammapicobirnaviridae* (Supplementary Figure 3), as well as two picobirnaviruses from vertebrate faecal samples (USE08169 Picobirnavirus sp. from pig faeces and QUS52969 mute swan feces associated picobirnavirus 3). However, most viruses from this genus were sampled in environmental or microbial samples, particularly aquatic sources.

Viruses in the genus *Deltapicobirnavirus* were predominantly sourced from terrestrial microbe communities and environments (Figure 3F, Supplementary Figure 4). The most recently diverged clade had a more even distribution of viruses sampled from aquatic and terrestrial environments and microbes (Supplementary Figure 4). Again, two picobirnaviruses sourced from animals were present in this genus - ND_241614 and ND_192065 - from cattle and sheep, respectively. However, these sequences were mined from animal rumen microbiome sequence data (Neri *et al*., 2022), thereby making *Deltapicobirnavirus* the only genus containing entirely microbe-associated or environmentally sourced viruses.

The remaining genera were considerably smaller. All but one of the 36 viruses falling into the genus *Epsilonpicobirnavirus* were sourced from vertebrate faeces, the majority of which were mammalian (cattle and porcine) (Figure 3C, Supplementary Figure 5). The genus *Zetapicobirnavirus* also featured a similar range of sampling sources, with the majority being cattle and porcine samples, along with one avian- and one primate-sourced picobirnavirus. This phylogeny included a notable divergent clade of five arthropod-associated zetapicobirnaviruses (Figure 3D, Supplementary Figure 6). Finally, despite only comprising 70 sequences, the genus *Etapicobirnavirus* had a broader “host” range across the two sister clades observed within this group. One predominantly comprised vertebrate etapicobirnaviruses, as well as one sequence identified in a termite that is likely associated with its bacterial symbionts (Figure 3G, Supplementary Figure 7). The other clade included another 15 etapicobirnaviruses identified in termite samples, with the basal lineages largely viruses sourced from microbial and aquatic environmental samples (Supplementary Figure 7).

### Accuracy of host assignments based on phylogenetic position

To visualise the extent of apparent cross-species transmission within the *Picobirnaviridae* based on assigned hosts/sample source, we estimated a phylogeny of only the animal-sourced picobirnaviruses (Figure 4A, T19 in Table 1) and a complete picobirnavirus phylogeny (Figure 4B, T9 in Table 1). Both trees were rooted by a set of 40 partitiviruses as they represent the most closely related family to the *Picobirnaviridae*. The diverse distribution of assigned host species across the phylogeny, as demonstrated in Figures 4A and 4B where tips are coloured by host, revealed a lack of consistent clustering by assigned host. Even at the order level within the *Mammalia*, closely related virus sequences had assigned hosts spanning diverse groups of mammals. Picobirnaviruses from birds did not form any large monophyletic groups, and instead clustered with species from cattle, pigs, felines, canines, primates, and marsupials (Figure 4A, C). This was also clearly observable within the predominantly animal-associated genera *Alphapicobirnavirus*, *Betapicobirnavirus*, *Epsilonpicobirnavirus*, and *Zetapicobirnavirus* (Figure 3A-D, Supplementary Figures 1-4). Although microbial/environmentally sourced viruses tended to group together in clades distinct from the animal-sourced picobirnaviruses, there were several invertebrate-, primate-, cattle-, pig-, and avian-associated species present in the environmental/microbial clades (Figure 4B, D). Notably, representatives from all sub-categories of microbial and environmental picobirnaviruses were present in animal-associated clades with the exception of those sourced from terrestrial microbes (Figure 4D). Rectangular, midpoint-rooted versions of the trees shown in Figure 4 with tip labels including accession numbers and virus names are shown in Supplementary Figures 8-11.

**Figure 4.**
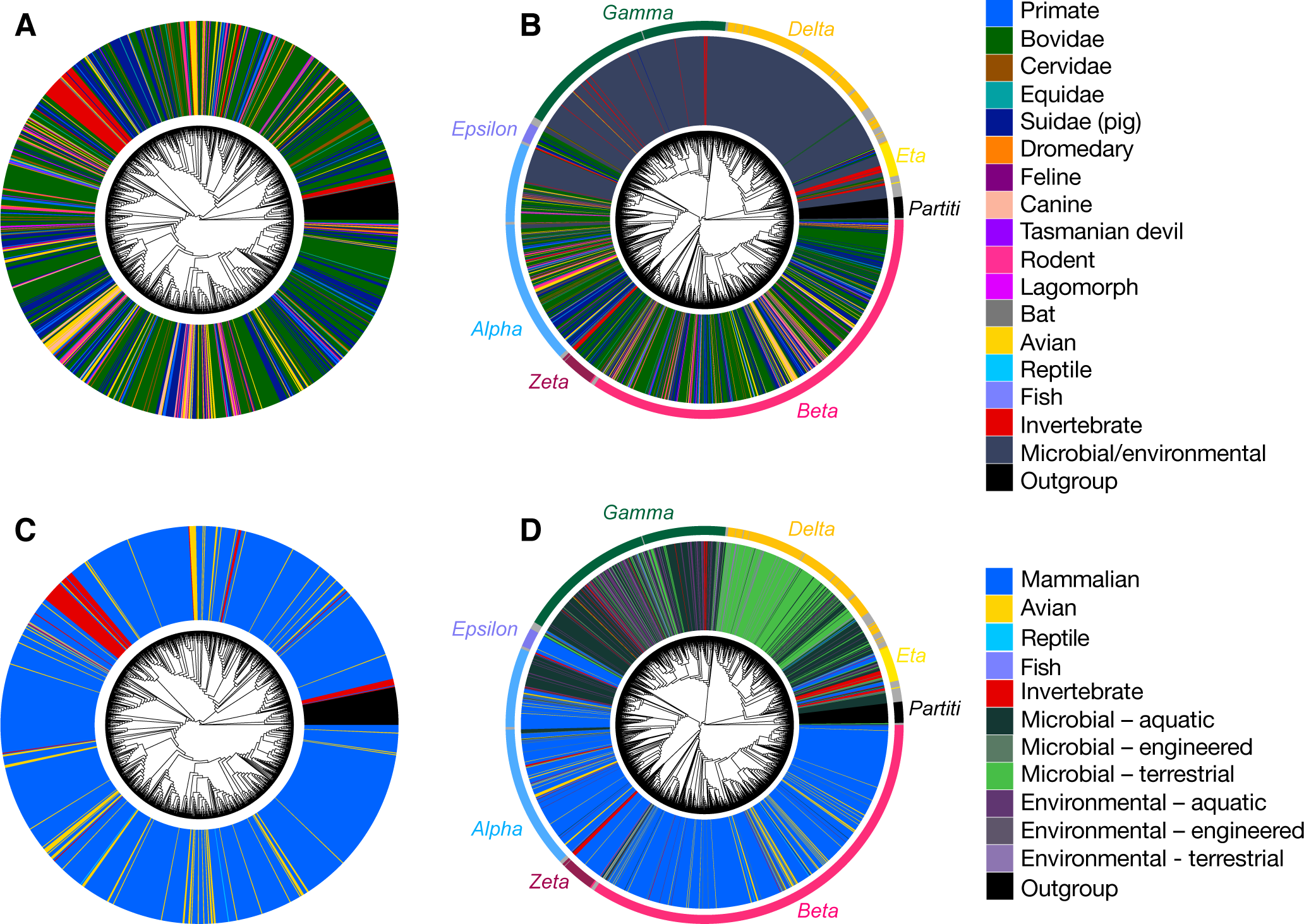
Maximum likelihood trees of *Picobirnaviridae* RdRp with tips coloured to represent assigned host/sampling environment. *Partitiviridae* RdRp sequences (n=40) were used to root all trees and are shown in black. The top two trees are coloured by animal host groups, including lower level Mammalia groupings, and are estimated on (A) *Picobirnaviridae* sequences with assumed animal associations and (B) all *Picobirnaviridae* sequences. The bottom two trees categorise animal hosts more broadly as ‘mammalian’, ‘avian’, ‘reptile’, ‘fish’, and ‘invertebrate’ and are estimated using (C) *Picobirnaviridae* sequences with assumed animal associations and (D) all *Picobirnaviridae* sequences with non-animal associated sequences grouped into ‘environmental’ or ‘microbial’ samples, both further classed into ‘terrestrial’, ‘aquatic’, and ‘engineered’ sampling sources. Scale bars for branch lengths are shown in rectangular versions of trees (Supplementary Figures 8-11).

A phylogeny of animal-associated picobirnaviruses (T20 in Table 1) was compared to a host phylogeny (Supplementary Figure 12, T21 in Table 1) using NELSI (Ho *et al*., 2015) to estimate the frequency of cross-species transmission (assuming assigned hosts are the true hosts) by calculating the nPH85 metric that describes the topological distance between the two phylogenies. The nPH85 distance between the unrooted animal-associated *Picobirnaviridae* tree and host taxa tree was 0.998 (Figure 5A), suggesting almost complete cross-species transmission and very little occurrence of co-divergence among the animal-associated members of this family. Similarly, the nPH85 distance for the genus *Alphapicobirnavirus* was 0.994, and 1 for both the *Betapicobirnavirus*, and *Epsilonpicobirnavirus.* The genus *Zetapicobirnavirus* had a comparatively higher, yet still low, level of co-divergence with an nPH85 distance of 0.921 (Figure 5A). eMPRess (Santichaivekin *et al*., 2021) was utilised to determine the respective likelihoods of co-divergence, cross-species transmission, duplication, or extinction in the reconciled virus-host co-phylogeny. The event likelihoods are shown in Figure 5B-F and reveal that cross-species transmission was far more frequent than any other event (68% of events, Figure 5B) for the entire family as well as for each individual genus (45-68% of events, Figure 5C-F). In contrast, only low levels of apparent virus-host co-divergence were observed, especially at the level of the entire *Picobirnaviriade* (1-23% of events, Figure 5B-F).

**Figure 5.**
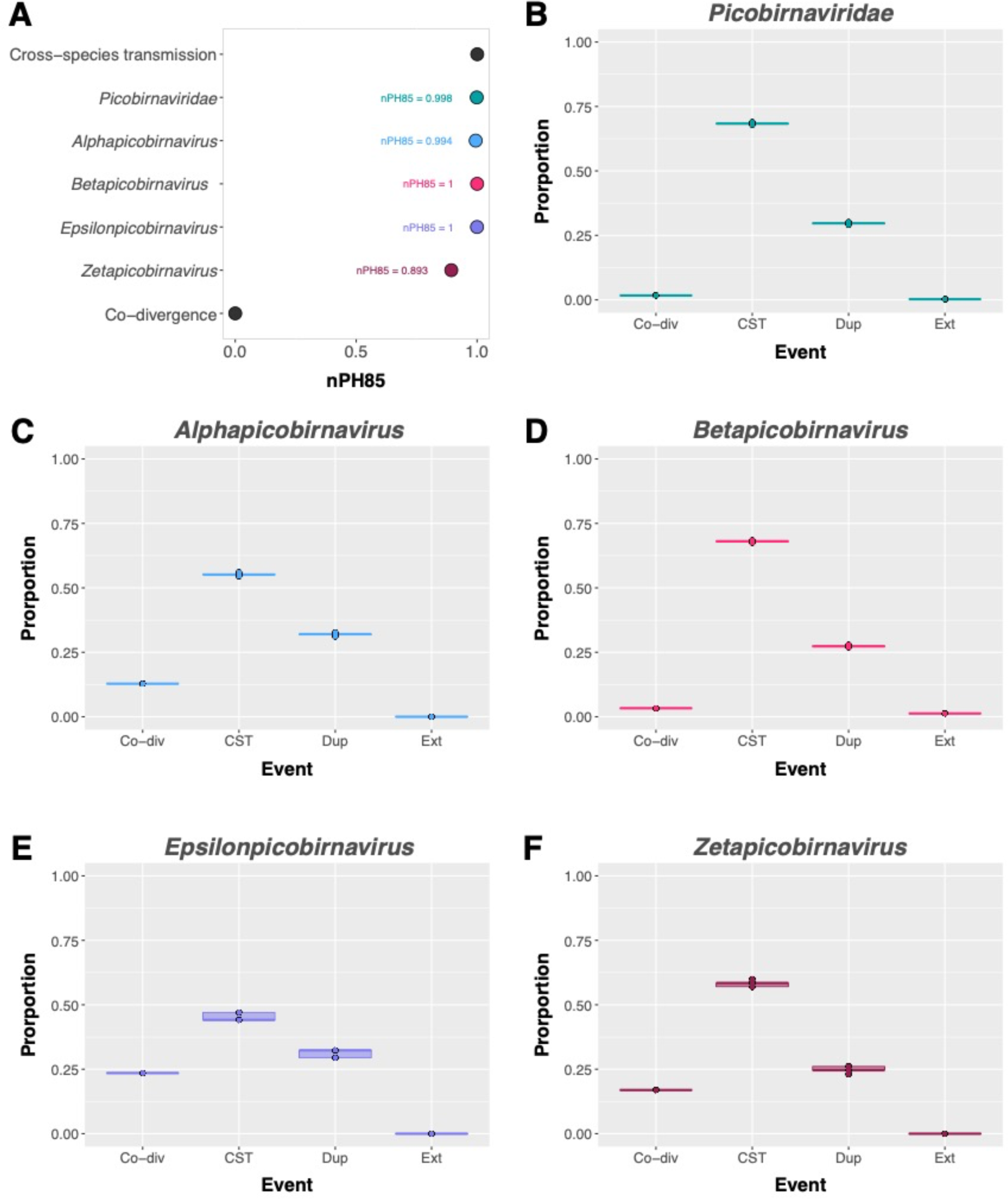
Levels of co-divergence of the *Picobirnaviridae* with associated animal hosts. (A) nPH85 metric calculated for the *Picobirnaviridae*, *Alphapicobirnavirus*, *Betapicobirnavirus, Epsilonpicobirnavirus,* and *Zetapicobirnavirus* using NELSI, representing the topological distance between unrooted virus and host trees. nPH85 = 0 indicates complete co-divergence and nPH85 = 1 indicates complete cross-species transmission. Event likelihoods of co-divergence (Co-div), cross-species transmission (CST), duplication (Dup), and extinction (Ext) as a proportion of all possible events in the reconciled co-phylogenies of animal-associated picobirnavirus sequences and their assigned hosts for (B) the family *Picobirnaviridae*, and the genera (C) *Alphapicobirnavirus*, (D) *Betapicobirnavirus*, (E) *Epsilonpicobirnavirus*, and (F) *Zetapicobirnavirus*. Event likelihoods were calculated in eMPRess with 100 replicates.

### Genetic evidence of *Picobirnaviridae* in non-animal hosts

We next searched for bacterial RBS motifs in a data set of 1,344 picobirnavirus, 1,040 levivirus, and 66 cystovirus genome segments. The detection of an ‘AGGAGG’ hexamer 14-24 nucleotides upstream of the start codon of an annotated ORF was considered to indicate the presence of an RBS motif. There were 722 annotated ORFs across the 360 picobirnavirus segment 1 nucleotide sequences and 993 annotated ORFs across the 984 picobirnavirus segment 2 sequences. The number of segment 1 and segment 2 ORFs preceded by an RBS motif was 431 (60%) and 665 (67%), respectively (Table 3). Strikingly, this was a higher proportion of RBS motif enrichment than observed in the confirmed RNA bacteriophage groups the *Leviviricetes* (210 of 3306 ORFs; 6.4%) and *Cystoviridae* (7 of 305 ORFs; 2.3%) (Table 3).

**Table 3.**
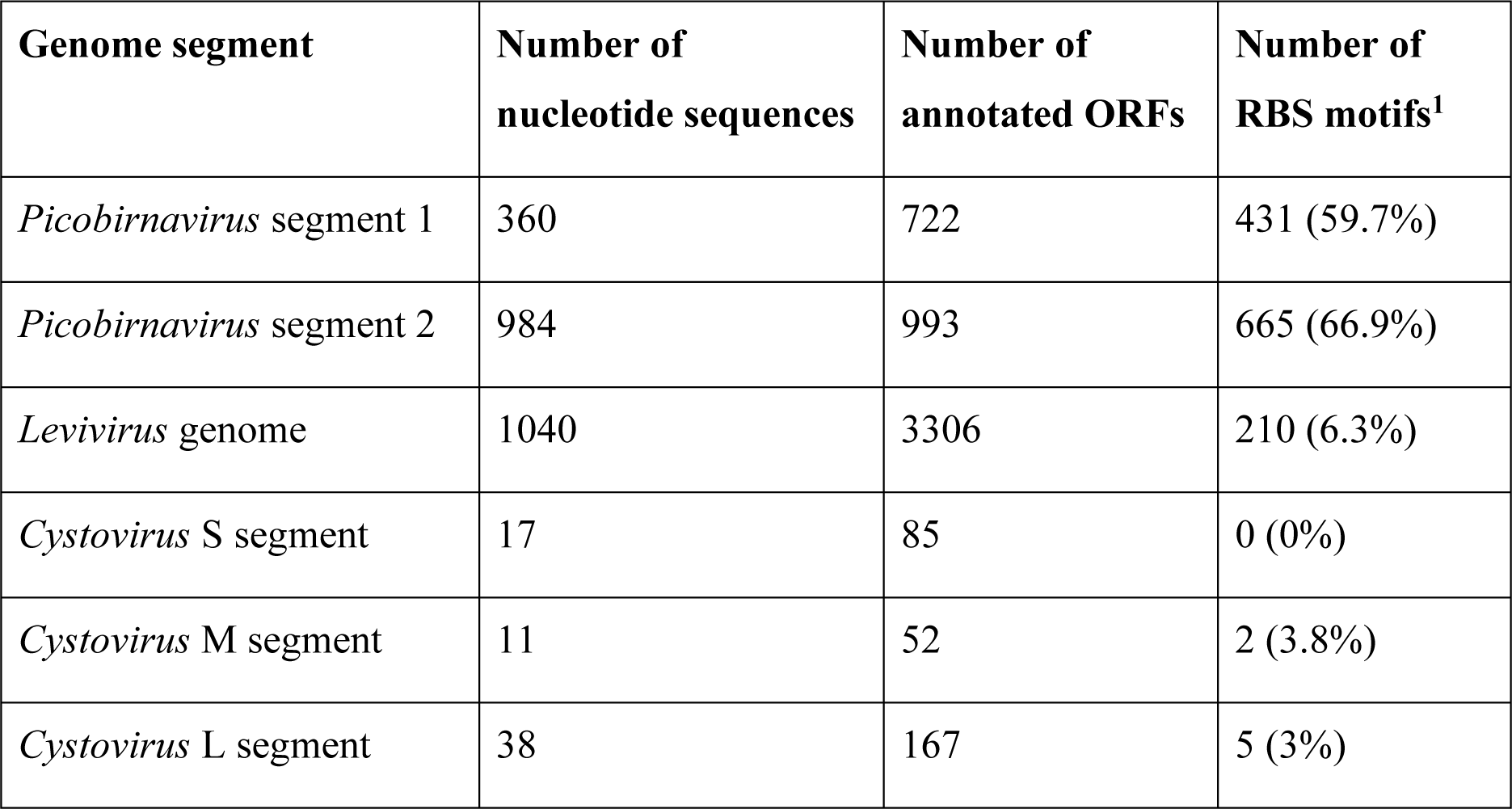
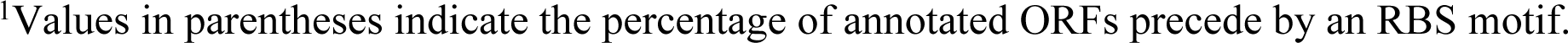
Genome segments and calculated proportions of annotated ORFs with the ‘AGGAGG’ RBS motif 14-24 nucleotides upstream of the start codon.

| Genome segment | Number of nucleotide sequences | Number of annotated ORFs | Number of RBS motifs<sup>1</sup> |
| --- | --- | --- | --- |
| *Picobirnavirus* segment 1 | 360 | 722 | 431 (59.7%) |
| *Picobirnavirus* segment 2 | 984 | 993 | 665 (66.9%) |
| *Levivirus* genome | 1040 | 3306 | 210 (6.3%) |
| *Cystovirus* S segment | 17 | 85 | 0 (0%) |
| *Cystovirus* M segment | 11 | 52 | 2 (3.8%) |
| *Cystovirus* L segment | 38 | 167 | 5 (3%) |

<sup>1</sup>Values in parentheses indicate the percentage of annotated ORFs precede by an RBS motif.

## DISCUSSION

The *Picobirnaviridae* are a family of RNA viruses traditionally associated with opportunistic gastroenteritis in humans and other animals that have dramatically increased in size and diversity in recent years due to metatranscriptomic sequencing. Despite this marked increase in diversity, picobirnaviruses remain largely unclassified beyond the family level, with only three divergent species comprising a single genus having been formally accepted by the ICTV (Delmas *et al*., 2019). The central aim of this study was to provide a taxonomic structure for this family by identifying distinct clades within the *Picobirnaviridae* that may represent genera (while noting that all such definitions are arbitrary). To this end, multiple outgroups and rooting schemes were utilised in an expansive phylogenetic analysis. From this, we identified seven clusters of sequences that consistently appeared across ten different phylogenies, which we propose be considered as distinct genera: *Alphapicobirnavirus*, *Betapicobirnavirus*, *Gammapicobirnavirus*, *Deltapicobirnavirus*, *Epsilonpicobirnavirus*, *Zetapicobirnavirus*, and *Etapicobirnavirus*. In addition, these seven proposed genera remained mostly intact and together formed a monophyletic, phylogenetically distinct group within the order *Durnavirales*. No genera shared greater than an average of 19% amino acid identity in the RdRp with any other genera, which is in line with the <24% amino acid identity shared between genera of the related *Partitiviridae* (Vainio *et al*., 2018).

Correctly assigning host organisms to novel virus species is a major limitation of the meta-transcriptomic approach to virus discovery. While the *Picobirnaviridae* have historically been associated with humans and other animals, there is mounting evidence that animals may not be the true hosts (Ghosh and Malik, 2021). As a consequence, we asked whether the *Picobirnaviridae* were a true animal-infecting family of RNA viruses, or one that infect microbiota themselves associated with animal hosts: i.e., that they are bacteriophage or other microbe-infecting viruses. The marked lack of topological congruence between phylogenies of the animal-associated picobirnaviruses and their apparent animal hosts indicates that the evolutionary history of the *Picobirnaviridae* has been characterised by extensive host jumping, with statistical tests revealing near complete cross-species transmission. Notably, the nPH85 distance for animal associated picobirnaviruses (0.998) is higher than the average for RNA virus families (0.95), and indeed higher than *Rhabdoviridae* (0.989), the family with the highest nPH85 distance of those analysed in Geoghegan *et al*. (2017). This supports previous suggestions that animal-associated picobirnaviruses do not cluster according to their “host” of sampling (Duraisamy *et al*., 2018; Woo *et al*., 2019; Mahar *et al*., 2020), although the current study uses a much larger data set. The complete lack of apparent host and viral phylogenetic congruence points toward an error in host assignment, rather than the less biologically feasible alternative of near complete cross-species transmission at a rate not seem in any other RNA virus family.

Picobirnaviruses have been detected in a large range of animal species, including those from terrestrial and aquatic habitats, invertebrates and vertebrates, and those with herbivorous, carnivorous, and omnivorous diets. This raises the question of how picobirnaviruses are able to overcome the geographical and biological barriers between animals with remarkably different habitats, lifestyles, and gastrointestinal systems, if they truly infect the hosts being sampled. Broad host ranges spanning vertebrates and invertebrates have been observed in several RNA virus orders, such as the *Articulavirales*, *Nidovirales*, *Reovirales*, and *Mononegevirales*, with the latter two also carrying fungi-infecting species. At a family level, the *Orthomyxoviridae*, *Rhabdoviridae*, and *Spinareoviridae* also infect a wide variety of hosts, including mammals, birds, insects and other arthropods, along with fish in the case of the *Orthomyxoviridae*, plants in the *Rhabdoviridae*, and fungi in the *Spinareoviridae*. There are, however, clear phylogenetic distinctions between genera infecting vastly different hosts, and host ranges are generally limited at a genus level. Genera with cross-phylum or cross-kingdom host ranges are typically characterised by well-established transmission routes between arthropod vectors and animal or plant hosts. For example, arboviruses in the genera *Orthoflavivirus*, *Alphavirus*, *Coltivirus*, and *Orbivirus* replicate in their vertebrate hosts as well as the arthropod vectors that transmit them (Huang *et al*., 2023). Similarly, plant-infecting genera in the families *Tospoviridae*, *Tymoviridae* (*Marafivirus*), *Rhabdoviridae*, and order *Reovirales* also replicate in their arthropod vectors (Gray and Banerjee, 1999). While it is plausible that members of the *Picobirnaviridae* are capable of extensive host-jumping, their remarkably wide host range raises the possibility that the picobirnaviruses are in fact associated with gut microflora or dietary components present in and excreted by these animals (Krishnamurthy and Wang, 2018). This would also explain why non-environmental picobirnaviruses are detected almost exclusively in animal faeces and are still unable to be cultured in any eukaryotic cell lines (Ghosh and Malik, 2021).

The presence of a bacterial ribosomal binding site motif is well-established in bacteriophage genomes. Krishnamurthy and Wang (2018) noted that the ‘AGGAGG’ RBS hexamer is enriched in the genomes of both RNA and DNA bacteriophage, and appears at much lower frequency in eukaryote-infecting virus genomes. In our expansive data set, we detected an RBS motif in 59-67% of ORFs in picobirnavirus genomes, such that this marker is substantially more enriched than in confirmed RNA bacteriophages *Leviviricetes* and *Cystoviridae*. This constitutes further evidence that the *Picobirnaviridae* indeed have bacterial hosts.

The host range of virus species within each of the seven proposed *Picobirnaviridae* genera appeared to comprise either predominantly animal-associated picobirnaviruses (*Alphapicobirnavirus*, *Betapicobirnavirus, Epsilonpicobirnavirus,* and *Zetapicobirnavirus*) or those identified in microbial or environmental samples (*Gammapicobirnavirus*, *Deltapicobirnavirus*, and *Etapicobirnavirus*). However, the four genera comprising predominantly animal-associated viruses did not group together to the exclusion of the three genera comprising mainly microbe/environmentally sourced viruses. As such, this broad host-associated clustering may reflect the different microbial community compositions in terrestrial, aquatic, and engineered environments compared to those of mammals, birds, and invertebrates. Picobirnaviruses may infect different microbial organisms that play roles in food webs linking vertebrates, invertebrates, and their diets and habitats. This would facilitate the extensive “host-jumping” observed within the animal-dominated genera. Hence, future metagenomic studies on the viromes of animal (particularly faecal) or environmental samples should also include community composition analyses of the microorganisms present. Determining if the microbial community composition of a sampled animal or environment is indeed driving the phylogenetic patterns of the *Picobirnaviridae* may elucidate the true host range of picobirnavirus genera.

## ACKNOWLEDGMENTS

This work was funded by a National Health and Medical Research Council (NHMRC) Investigator award (GNT201719) to ECH and by AIR@InnoHK administered by the Innovation and Technology Commission, Hong Kong Special Administrative Region, China.

## AUTHOR CONTRIBUTIONS

SS, ECH and JEM conceptualised this study. SS collated and analysed the sequence data. SS wrote the original manuscript draft. SS, ECH and JEM edited and revised the manuscript. ECH and JEM supervised the project.

