## Supplementary Figures for "Genomic and phylogenetic features of the *Picobirnaviridae* suggest microbial rather than animal hosts"

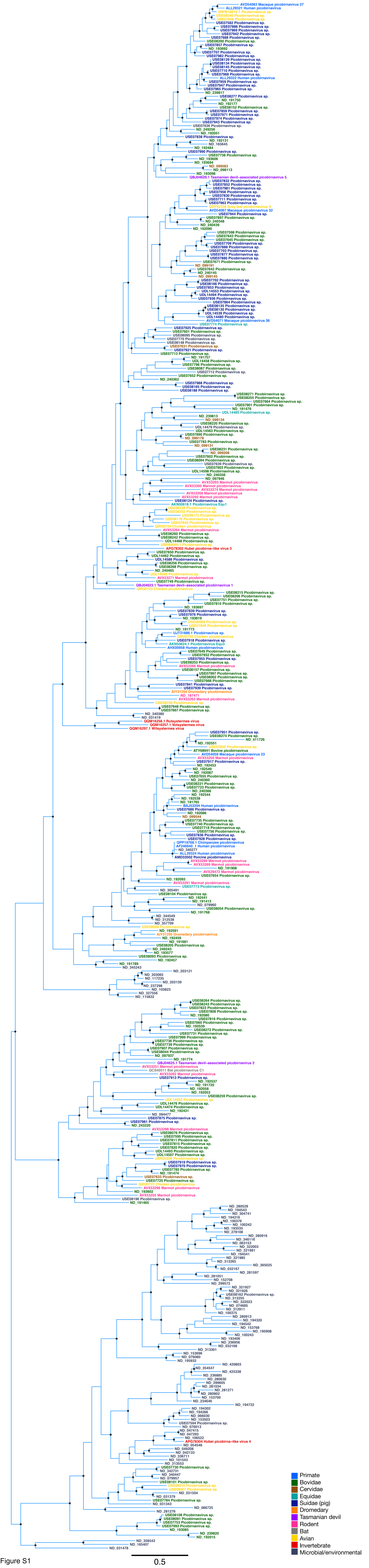
Primate
Bovidae
Cervidae
Equidae
Suidae (pig)
Dromedary
Tasmanian devil
Rodent
Bat
Avian
Invertebrate
Microbial/environmental
0.5

Figure S1

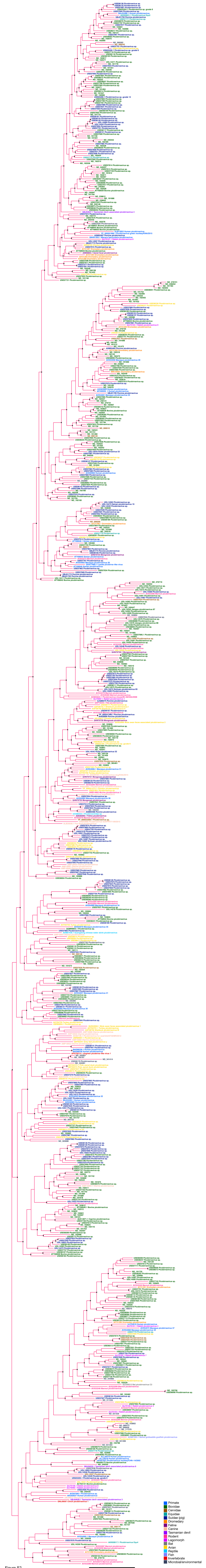
Primate
Bovidae
Cervidae
Equidae
Suidae (pig)
Dromedary
Feline
Canine
Tasmanian devil
Rodent
Lagomorph
Bat
Avian
Reptile
Fish
Invertebrate
Microbial/environmental
0.5

Figure S2

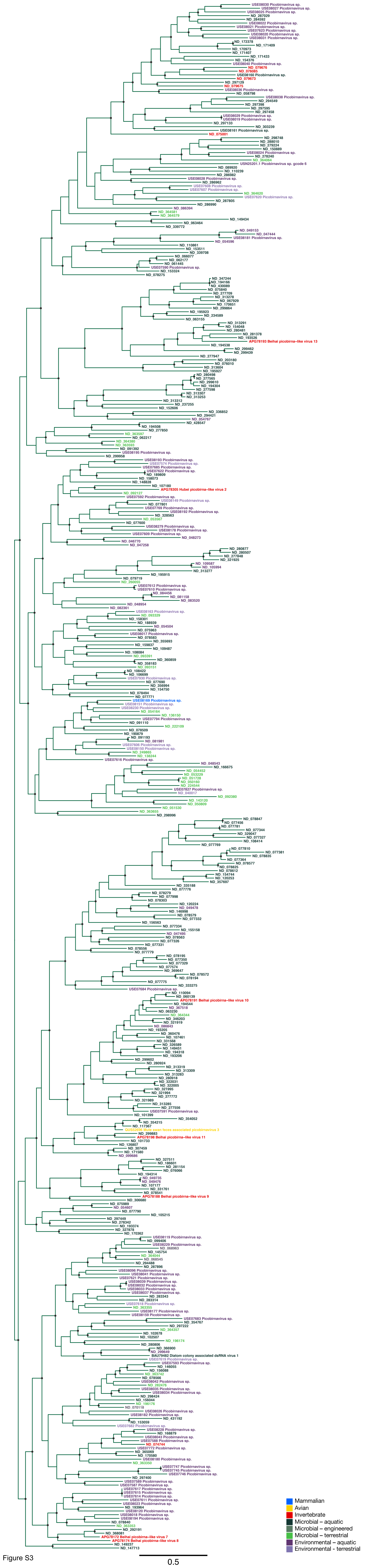
Mammalian
Avian
Invertebrate
Microbial – aquatic
Microbial – engineered
Microbial – terrestrial
Environmental – aquatic
Environmental – terrestrial
0.5

Figure S3

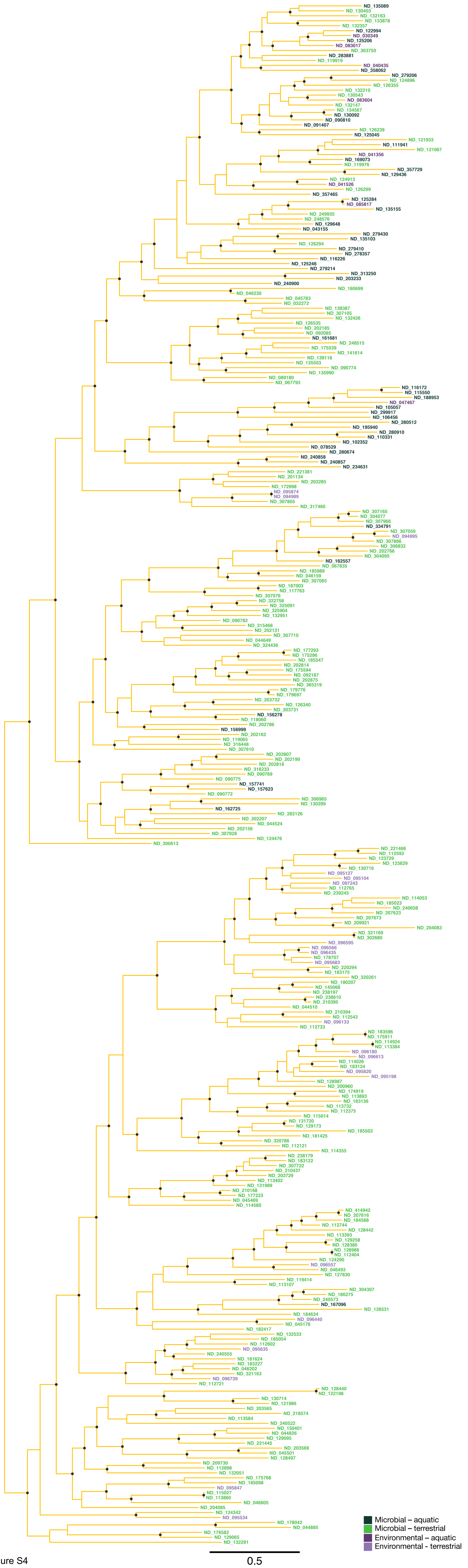

Microbial – aquatic
Microbial – terrestrial
Environmental – aquatic
Environmental - terrestrial
0.5


Figure S4

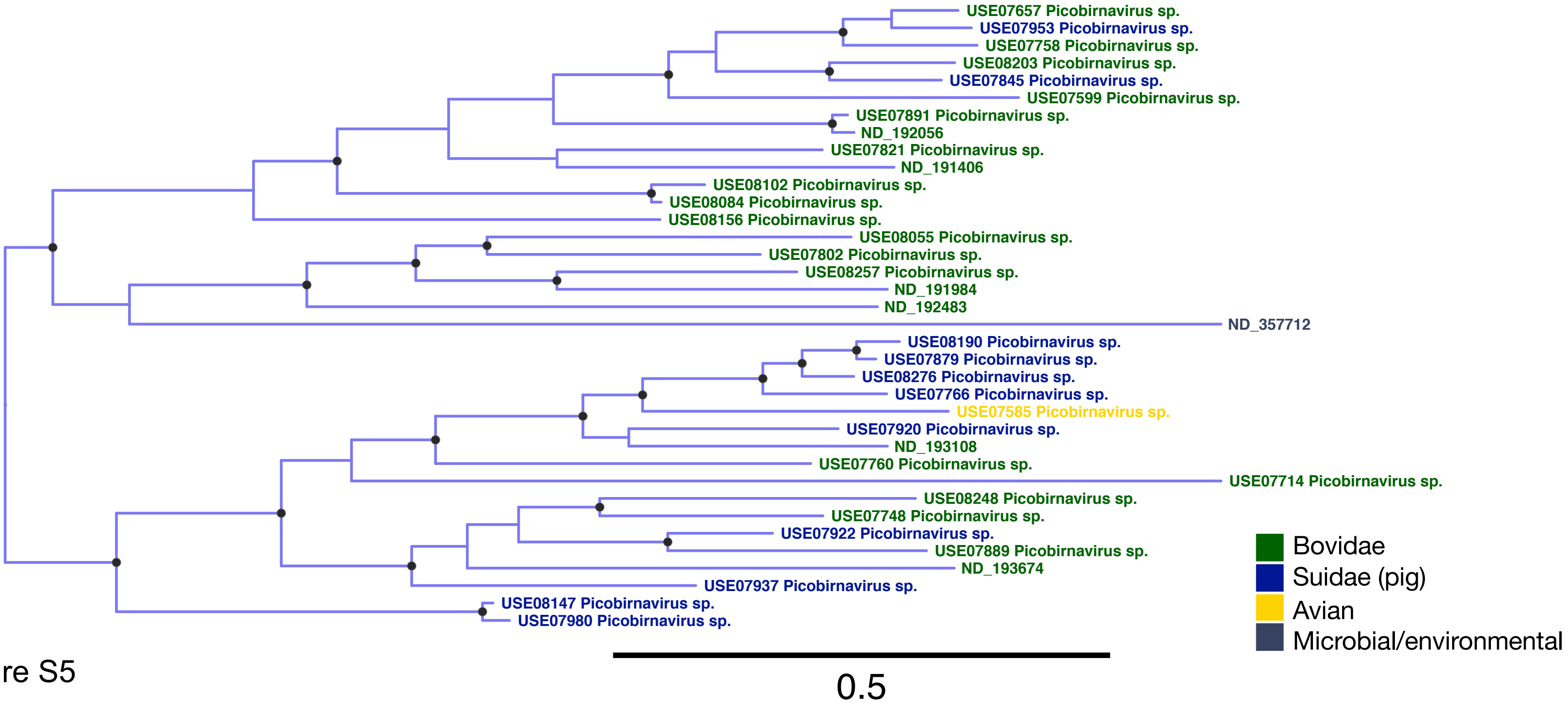
USE07657 Picobirnavirus sp.
USE07953 Picobirnavirus sp.
USE07758 Picobirnavirus sp.
USE08203 Picobirnavirus sp.
USE07845 Picobirnavirus sp.
USE07599 Picobirnavirus sp.
USE07891 Picobirnavirus sp.
ND_192056
USE07821 Picobirnavirus sp.
ND_191406
USE08102 Picobirnavirus sp.
USE08084 Picobirnavirus sp.
USE08156 Picobirnavirus sp.
USE08055 Picobirnavirus sp.
USE07802 Picobirnavirus sp.
USE08257 Picobirnavirus sp.
ND_191984
ND_192483
ND_357712
USE08190 Picobirnavirus sp.
USE07879 Picobirnavirus sp.
USE08276 Picobirnavirus sp.
USE07766 Picobirnavirus sp.
USE07585 Picobirnavirus sp.
USE07920 Picobirnavirus sp.
ND_193108
USE07760 Picobirnavirus sp.
USE07714 Picobirnavirus sp.
USE08248 Picobirnavirus sp.
USE07748 Picobirnavirus sp.
USE07922 Picobirnavirus sp.
USE07889 Picobirnavirus sp.
ND_193674
USE07937 Picobirnavirus sp.
USE08147 Picobirnavirus sp.
USE07980 Picobirnavirus sp.
Bovidae
Suidae (pig)
Avian
Microbial/environmental
0.5

Figure S5

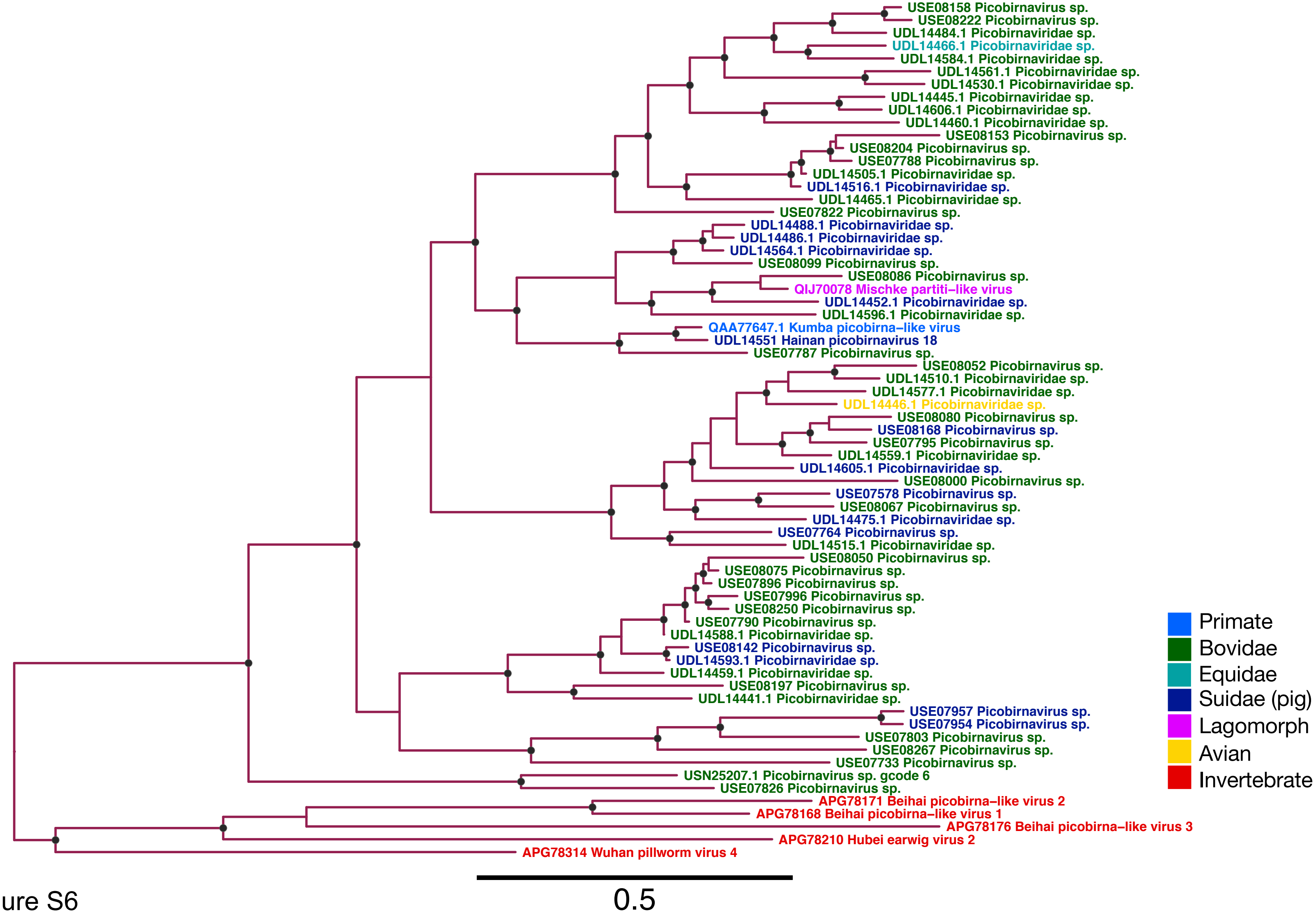
USE08158 Picobirnavirus sp.
USE08222 Picobirnavirus sp.
UDL14484.1 Picobirnaviridae sp.
UDL14466.1 Picobirnaviridae sp.
UDL14584.1 Picobirnaviridae sp.
UDL14561.1 Picobirnaviridae sp.
UDL14530.1 Picobirnaviridae sp.
UDL14445.1 Picobirnaviridae sp.
UDL14606.1 Picobirnaviridae sp.
UDL14460.1 Picobirnaviridae sp.
USE08153 Picobirnavirus sp.
USE08204 Picobirnavirus sp.
USE07788 Picobirnavirus sp.
UDL14505.1 Picobirnaviridae sp.
UDL14516.1 Picobirnaviridae sp.
UDL14465.1 Picobirnaviridae sp.
USE07822 Picobirnavirus sp.
UDL14488.1 Picobirnaviridae sp.
UDL14486.1 Picobirnaviridae sp.
UDL14564.1 Picobirnaviridae sp.
USE08099 Picobirnavirus sp.
USE08086 Picobirnavirus sp.
QIJ70078 Mischke partiti−like virus
UDL14452.1 Picobirnaviridae sp.
UDL14596.1 Picobirnaviridae sp.
QAA77647.1 Kumba picobirna−like virus
UDL14551 Hainan picobirnavirus 18
USE07787 Picobirnavirus sp.
USE08052 Picobirnavirus sp.
UDL14510.1 Picobirnaviridae sp.
UDL14577.1 Picobirnaviridae sp.
UDL14446.1 Picobirnaviridae sp.
USE08080 Picobirnavirus sp.
USE08168 Picobirnavirus sp.
USE07795 Picobirnavirus sp.
UDL14559.1 Picobirnaviridae sp.
UDL14605.1 Picobirnaviridae sp.
USE08000 Picobirnavirus sp.
USE07578 Picobirnavirus sp.
USE08067 Picobirnavirus sp.
UDL14475.1 Picobirnaviridae sp.
USE07764 Picobirnavirus sp.
UDL14515.1 Picobirnaviridae sp.
USE08050 Picobirnavirus sp.
USE08075 Picobirnavirus sp.
USE07896 Picobirnavirus sp.
USE07996 Picobirnavirus sp.
USE08250 Picobirnavirus sp.
USE07790 Picobirnavirus sp.
UDL14588.1 Picobirnaviridae sp.
USE08142 Picobirnavirus sp.
UDL14593.1 Picobirnaviridae sp.
UDL14459.1 Picobirnaviridae sp.
USE08197 Picobirnavirus sp.
UDL14441.1 Picobirnaviridae sp.
USE07957 Picobirnavirus sp.
USE07954 Picobirnavirus sp.
USE07803 Picobirnavirus sp.
USE08267 Picobirnavirus sp.
USE07733 Picobirnavirus sp.
USN25207.1 Picobirnavirus sp. gcode 6
USE07826 Picobirnavirus sp.
APG78171 Beihai picobirna−like virus 2
APG78168 Beihai picobirna−like virus 1
APG78176 Beihai picobirna−like virus 3
APG78210 Hubei earwig virus 2
APG78314 Wuhan pillworm virus 4
Primate
Bovidae
Equidae
Suidae (pig)
Lagomorph
Avian
Invertebrate
0.5

Figure S6

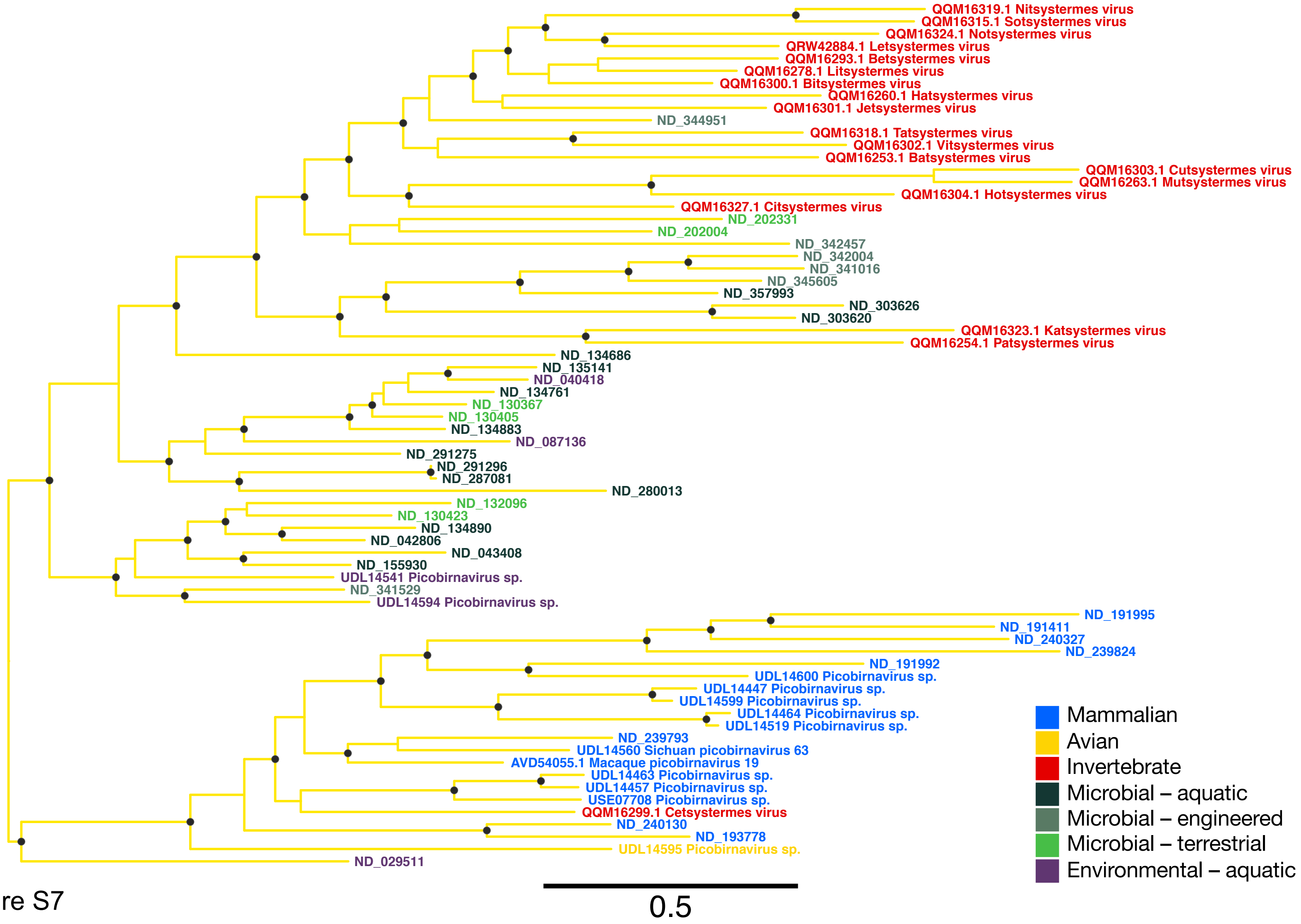
QQM16319.1 Nitsystermes virus
QQM16315.1 Sotsystermes virus
QQM16324.1 Notsystermes virus
QRW42884.1 Letsystermes virus
QQM16293.1 Betsystermes virus
QQM16278.1 Litsystermes virus
QQM16300.1 Bitsystermes virus
QQM16260.1 Hatsystermes virus
QQM16301.1 Jetsystermes virus
ND_344951
QQM16318.1 Tatsystermes virus
QQM16302.1 Vitsystermes virus
QQM16253.1 Batsystermes virus
QQM16303.1 Cutsystermes virus
QQM16263.1 Mutsystermes virus
QQM16304.1 Hotsystermes virus
QQM16327.1 Citsystermes virus
ND_202331
ND_202004
ND_342457
ND_342004
ND_341016
ND_345605
ND_357993
ND_303626
ND_303620
QQM16323.1 Katsystermes virus
QQM16254.1 Patsystermes virus
ND_134686
ND_135141
ND_040418
ND_134761
ND_130367
ND_130405
ND_134883
ND_087136
ND_291275
ND_291296
ND_287081
ND_280013
ND_132096
ND_130423
ND_134890
ND_042806
ND_043408
ND_155930
UDL14541 Picobirnavirus sp.
ND_341529
UDL14594 Picobirnavirus sp.
ND_191995
ND_191411
ND_240327
ND_239824
ND_191992
UDL14600 Picobirnavirus sp.
UDL14447 Picobirnavirus sp.
UDL14599 Picobirnavirus sp.
UDL14464 Picobirnavirus sp.
UDL14519 Picobirnavirus sp.
ND_239793
UDL14560 Sichuan picobirnavirus 63
AVD54055.1 Macaque picobirnavirus 19
UDL14463 Picobirnavirus sp.
UDL14457 Picobirnavirus sp.
USE07708 Picobirnavirus sp.
QQM16299.1 Cetsystermes virus
ND_240130
ND_193778
UDL14595 Picobirnavirus sp.
ND_029511
0.5
Mammalian
Avian
Invertebrate
Microbial – aquatic
Microbial – engineered
Microbial – terrestrial
Environmental – aquatic

Figure S7

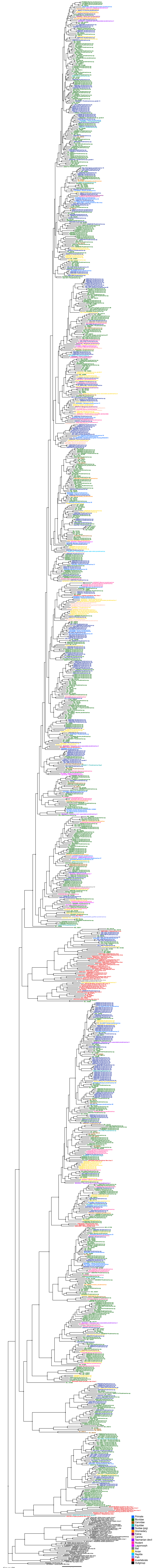

Primate
Bovidae
Cervidae
Equidae
Suidae (pig)
Dromedary
Feline
Canine
Tasmanian devil
Rodent
Lagomorph
Bat
Avian
Reptile
Fish
Invertebrate
Outgroup
0.5

Figure S8

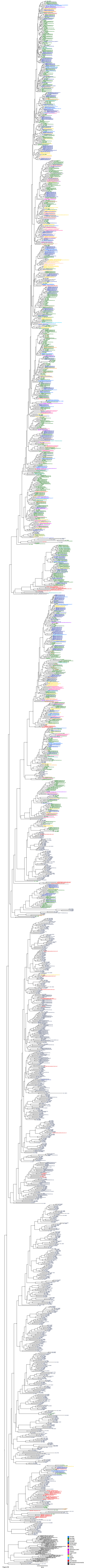
0.5
Primate
Bovidae
Cervidae
Equidae
Suidae (pig)
Dromedary
Feline
Canine
Tasmanian devil
Rodent
Lagomorph
Bat
Avian
Reptile
Fish
Invertebrate
Microbial/environmental
Outgroup

Figure S9

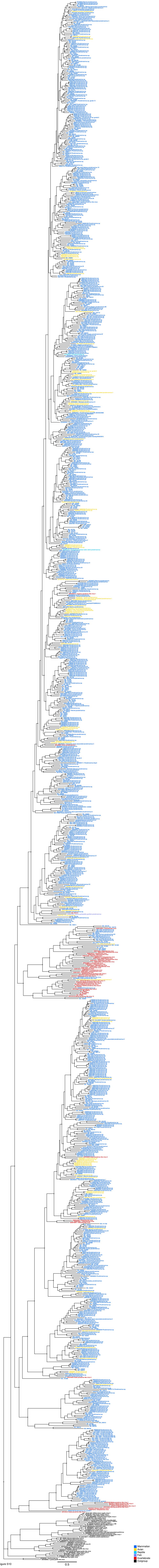


Figure S10

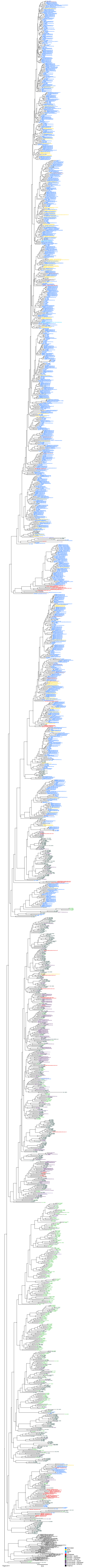
Mammalian
Avian
Reptile
Fish
Invertebrate
Microbial – aquatic
Microbial – engineered
Microbial – terrestrial
Environmental – aquatic
Environmental – engineered
Environmental - terrestrial
Outgroup
0.5

Figure S11

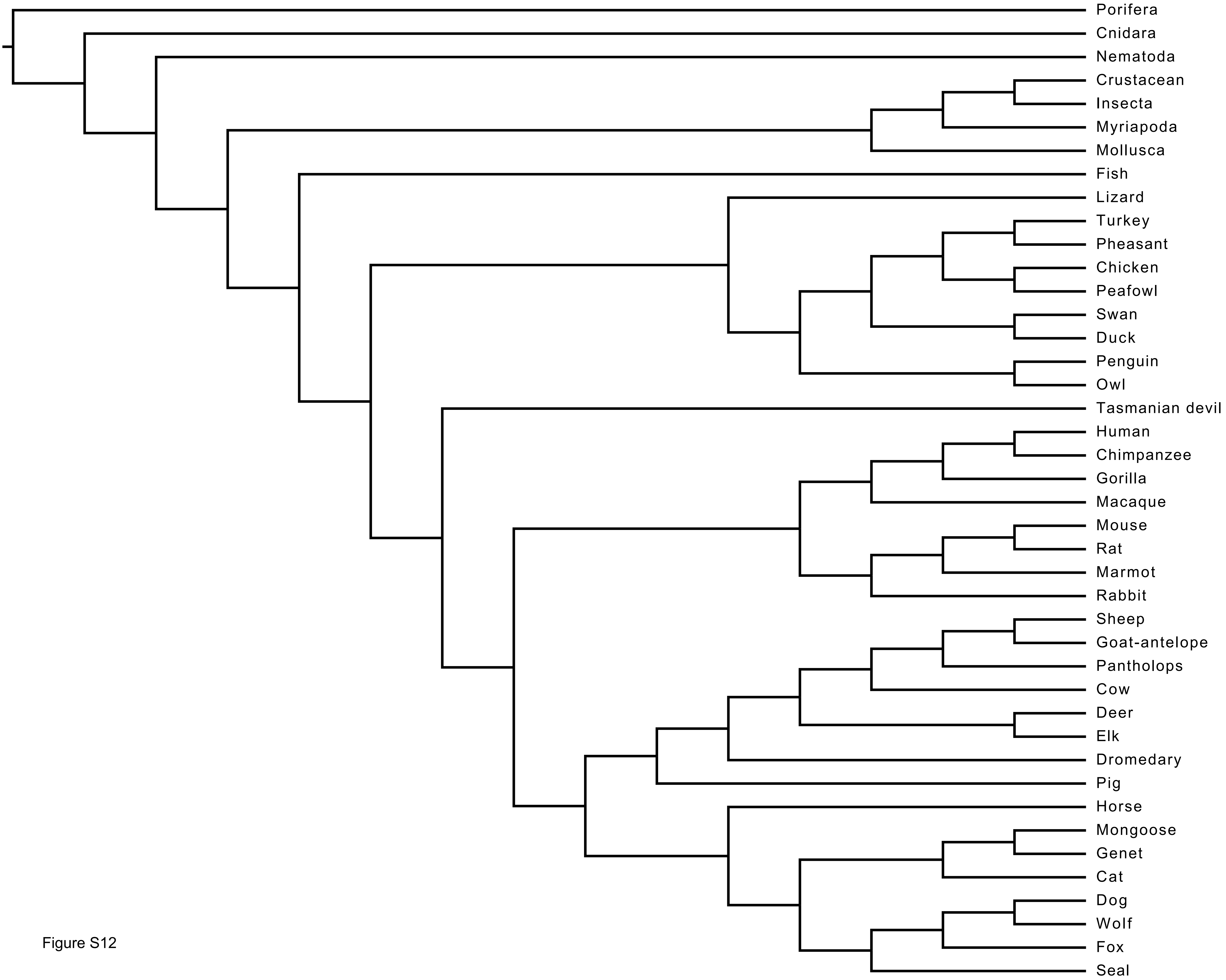
Porifera
Cnidara
Nematoda
Crustacean
Insecta
Myriapoda
Mollusca
Fish
Lizard
Turkey
Pheasant
Chicken
Peafowl
Swan
Duck
Penguin
Owl
Tasmanian devil
Human
Chimpanzee
Gorilla
Macaque
Mouse
Rat
Marmot
Rabbit
Sheep
Goat-antelope
Pantholops
Cow
Deer
Elk
Dromedary
Pig
Horse
Mongoose
Genet
Cat
Dog
Wolf
Fox
Seal

Figure S12
